# Central lipid sensing tunes accumbal D2R-neuron activity in appetitive behaviours

**DOI:** 10.64898/2026.09.21.753293

**Authors:** Guangping Li, Julien Castel, Anthony Ansoult, Benoit Bertrand, Giuseppe Gangarossa, Serge Luquet

## Abstract

Energy homeostasis and motivated behaviours are tightly interconnected processes coordinated by peripheral metabolic signals and central reward circuits. Among these signals, circulating triglycerides (TG) have emerged as neuromodulators of mesolimbic dopamine functions and reward processing. However, whether TG directly regulate the activity of defined neuronal populations within the nucleus accumbens (NAc) during appetitive behaviours remains unknown. Here, we investigated how central TG availability shapes the *in vivo* activity of NAc dopamine D2 receptor-expressing spiny projection neurons (D2R-SPNs) across distinct nutritional and metabolic states. Using *in vivo* fiber photometry in *Drd2*-Cre mice, we monitored D2R-SPN Ca^2+^ dynamics during appetitive Pavlovian conditioning and refeeding. D2R-SPNs progressively developed robust responses to a reward-predictive conditioned stimulus (CS^+^) over the course of learning, while their activity during reward consumption remained stable. Acute elevation of central TG levels through carotid infusion did not alter D2R-SPN activity or behavioural performance in lean chow-fed mice, irrespective of whether they were fed or food-restricted. In contrast, in mice exposed to a high-fat diet (HFD), central TG delivery suppressed cue-evoked D2R-SPN activity in food-restricted animals without affecting reward consumption or behavioural outputs. Likewise, TG significantly reduced refeeding-evoked D2R-SPN activation in fasted HFD-mice, while leaving novelty-induced neuronal responses unchanged.

Together, these findings demonstrate that obesogenic conditions reveal a metabolic state-dependent sensitivity of accumbal D2R-SPNs to circulating lipids. This work identifies TG as context-dependent modulators of D2R-SPN function and uncovers a mechanism through which dietary history interacts with current metabolic state to reshape the neural encoding of reward-predictive cues and food-directed behaviours.

## Introduction

Energy homeostasis and motivated behaviours are tightly interconnected processes that enable organisms to adapt feeding strategies to fluctuating energetic demands. Beyond their classical role as energy substrates, circulating lipids, central lipid deposit and lipid metabolism are increasingly recognized as key biochemical actors capable of influencing neural properties and circuits ^1–6^, some of which are involved in reward processing and motivation ^2,7–9^. Among circulating lipids, triglycerides (TG) represent a major metabolic signal whose circulating levels dynamically fluctuate according to nutritional states, increasing after food intake and during obesogenic conditions ^10^. Elevated circulating TG levels are commonly associated with obesity and metabolic syndrome, both of which are characterized by profound alterations in reward-related behaviours and mesolimbic dopamine (DA) signaling in both humans and rodents ^11–19^. Yet, the mechanisms through which TG influence neuronal activity within reward circuits during appetitive behaviours remain largely unresolved.

The nucleus accumbens (NAc), a central component of the mesolimbic DA system, integrates peripheral metabolic signals with motivational and reward-related processes. DA transmission within the NAc is critically involved in appetitive learning, cue-reward associations, and food-seeking behaviours ^20–23^. In particular, DA D2 receptor-expressing spiny projection neurons (D2R-SPNs) play a pivotal role in behavioural adaptation, motivational control, and action selection. In fact, alterations in accumbal D2R signaling have been consistently associated with obesity, compulsive eating, and addiction-like phenotypes (for review ^24^), suggesting that metabolic disturbances profoundly reshape the functional organization of mesolimbic circuits.

Accumulating evidence further indicates that dietary lipids directly interact with mesolimbic structures to modulate reward processing. Notably, we have shown that central delivery of TG suppresses motivation for palatable food and alters reward-seeking behaviours, while disruption of the TG-processing enzyme lipoprotein-lipase (LPL) in the NAc enhances food preference and motivational drive ^2,25^. These findings support the concept that TG act not only as metabolic substrates but also as “neuromodulatory” signals ^2,3,5^ capable of regulating mesolimbic functions ^7^. However, whether TG specifically influence the activity of identified neuronal populations within the NAc during appetitive behaviours remains unknown.

Metabolic state itself exerts a strong influence on mesolimbic DA signaling. Fasting enhances motivational responses to food-related cues and sensitizes reward circuits, whereas refeeding suppresses food-directed behaviours and reorganizes DA-dependent neural activity. Food restriction and weight loss are associated with marked alterations in extracellular DA levels within the NAc ^21,22,26–29^, highlighting the sensitivity of mesolimbic pathways to energy states. These observations suggest that peripheral metabolic signals, including TG, may dynamically tune accumbal neuronal activity according to the organism’s current energetic needs.

In parallel, short-term exposure to obesogenic diets is increasingly recognized as sufficient to induce early functional alterations in reward circuitry before the development of overt obesity. High-fat high-sucrose (HFHS) diets rapidly modify DA signaling, reward sensitivity, and food-seeking behaviours ^11,30–33^, thereby potentially altering the integration of peripheral lipids-derived signals by accumbal neurons. Therefore, understanding how metabolic states and dietary history interact to regulate specific neuronal populations within the NAc represents a critical challenge for elucidating the neural bases of maladaptive feeding behaviours.

Here, we investigated whether centrally delivered TG tune the activity of accumbal D2R-SPNs during appetitive behaviour in a metabolic state-dependent manner. To address this question, we combined an appetitive Pavlovian conditioning paradigm with fasting and refeeding challenges in lean mice as well as in mice exposed short-term to HFD. Using *in vivo* monitoring of D2R-SPN activity during cue-reward association and food-directed behaviours, we examined how physiological and diet-induced fluctuations in TG levels shape accumbal neuronal dynamics under fed and fasted conditions. Our findings identify triglycerides as key metabolic modulators of accumbal D2R-SPN activity and reveal a mechanism through which peripheral lipid signals adapt reward-related neural processing to nutritional states and dietary environment.

## Material and Methods

### Animals

All animal experiments were performed with approval of the Animal Care Committee of the Université Paris Cité (Apafis #29892). 8-12 weeks old male *Drd2*-Cre mice (STOCK Tg(Drd2-cre)ER44Gsat/Mmucd) were individually housed in a room maintained at 22 ± 1°C with a light period from 7h00 to 19h00. Regular chow diet (3.24 kcal/g, reference SAFE® A04, Augy, France), high-fat diet (HFD, Research Diets, Cat #D12492, 5.24 kcal/g) and water were provided ad libitum unless otherwise stated. All procedures were designed to minimize animal suffering and reduce the number of animals used. Behavioural tests occurred during the light phase, notably between 9 AM and 5 PM.

### Behavioural experiments

#### Appetitive Pavlovian conditioning

The appetitive pavlovian conditioning protocol was adapted from a previous study ^34^. Briefly, first, mice were placed in the operant chamber for environmental habituation to minimize stress and neophobia. Next, food-restricted (10% reduction from their initial body weight) or *ad libitum* fed mice were placed in the operant chamber for 20-min sessions. During each session, a drop of sweetened condensed milk was delivered 3 s after the onset of a sound (65 dB, 3000 Hz, 7 s) and illumination of a diode (10 s). The inter-trial interval (ITI) was 45 s following food-reward consumption. The food-reward had to be consumed by the animal in order for a new reward to be delivered. Animals were trained for 20 sessions. Licking patterns, including the number of licks and lick bursts, were analysed as a proxy for food-reward consumption.

#### Fasting and refeeding

Mice were fasted overnight for 14 hours and, depending on their experimental group, were subsequently exposed to either a chow diet (CD) or a high-fat diet (HFD). Food intake was measured at 30-min intervals over a 3h period (180 min).

#### Amphetamine-induced D2R-SPN inhibition

Amphetamine (2 mg/kg, i.p., Tocris, #2813) was administered (i.p.) to *Drd2*-Cre mice to measure Ca^2+^ transients (fiber photometry) in D2R-SPNs.

### Catheter implantation and infusion procedures

Surgery and central perfusions were carried out as previously described ^2,25^. Mice were anesthetized with isoflurane (3.5% for induction, 1.5% for maintenance) and received 10 mg/kg intraperitoneal injection (i.p.) of Buprécare® (Buprenorphine 0.3 mg) diluted 1/100 in NaCl 0.9% and 10 mg/kg of Ketofen® (Ketoprofen 100 mg) diluted 1/100 in NaCl 0.9%. Home-made catheters were inserted in the left carotid artery towards the brain. Catheters clotting was prevented through regular flushing with small volumes of NaCl 0.9%. Infusions started after a recovery period of 7 days by connecting catheters to a swivelling infusion device allowing animals to move freely and access water and food. Mice were habituated to the infusion device for two consecutive days. After the pavlovian training sessions, mice received NaCl 0.9% (Sal mice) or TG emulsion (TG mice) (Intralipid™ 20%) at a rate of 0.5 μl/min for 90 minutes.

### Viral production

pAAV.Syn.Flex.GCaMP6f.WPRE.SV40 (titer ≥ 1×10^13^ vg/ml, working dilution 1:5) was a gift from Douglas Kim (Addgene viral prep #100833-AAV9; https://www.addgene.org/100833/; RRID:Addgene_100833).

### Stereotaxic procedures

Mice were anaesthetized with isoflurane and received 10 mg/kg intraperitoneal injection (i.p.) of Buprécare® (Buprenorphine 0.3 mg) diluted 1/100 in NaCl 0.9% and 10 mg/kg of Ketofen® (Ketoprofen 100 mg) diluted 1/100 in NaCl 0.9%, and placed on a stereotactic frame (Model 940, David Kopf Instruments, California). The virus (0.3 μl) was injected unilaterally (fiber photometry) into the nucleus accumbens (NAc) (L=+/−1; AP=+1; V=−4.2, in mm) at a rate of 0.1 μl/min. The injection needle was carefully removed after 5 minutes waiting at the injection site and 2 minutes waiting half way to the top. Optical fiber for Ca^2+^ imaging into the NAc was implanted 100 μm above the viral injection site.

### Fiber photometry and data analysis

A chronically implantable cannula (Doric Lenses, Québec, Canada) composed of a bare optical fiber (400 μm core, 0.48 N.A.) and a fiber ferrule was implanted 100 μm above the location of the viral injection site in the nucleus accumbens (NAc: L=+/−1; AP=+1; V=−4.2, in mm). The fiber was fixed onto the skull using dental cement (Super-Bond C&B, Sun Medical). Real time fluorescence emitted from the calcium sensor GCaMP6f expressed by D2R-neurons was recorded using fiber photometry as previously described in the literature ^2,35,36^. Fluorescence was collected in the NAc using a single optical fiber for both delivery of excitation light streams and collection of emitted fluorescence.

The fiber photometry setup used 2 light emitting LEDs: 405 nm LED sinusoidally modulated at 330 Hz and a 465 nm LED sinusoidally modulated at 533 Hz (Doric Lenses) merged in a FMC4 MiniCube (Doric Lenses) that combines the 2 wavelengths excitation light streams and separate them from the emission light. The MiniCube was connected to a Fiberoptic rotary joint (Doric Lenses) connected to the cannula. A RZ5P lock-in digital processor controlled by the Synapse software (Tucker-Davis Technologies, TDT, USA), commanded the voltage signal sent to the emitting LEDs via the LED driver (Doric Lenses). The light power before entering the implanted cannula was measured with a power meter (PM100USB, Thorlabs) before the beginning of each recording session. The irradiance was ∼9 mW/cm^2^. The fluorescence emitted by the GCaMP6f activation in response to light excitation was collected by a femtowatt photoreceiver module (Doric Lenses) through the same fiber patch cord. The signal was then received by the RZ5P processor (TDT). On-line real time demodulation of the fluorescence due to the 405nm and the 465 nm excitations was performed by the Synapse software (TDT). A camera was synchronized with the recording using the Synapse software. Signals were exported and analysed using pMAT ^37^.

### Tissue preparation and immunofluorescence

Mice were rapidly anaesthetized with pentobarbital (500 mg/kg, i.p., Sanofi-Aventis, France) and transcardially perfused with 4% (weight/vol.) paraformaldehyde in 0.1 M sodium phosphate buffer (pH 7.5). Brains were post-fixed overnight in the same solution and stored at 4°C. 40 μm-thick sections were cut with a vibratome (Leica VT1000S, France), stored at −20 °C in a solution containing 30% ethylene glycol, 30% glycerol and 0.1 M sodium phosphate buffer. GCaMP6f-positive neurons were visualized without amplification and slices were counterstained with DAPI.

Acquisitions were performed with a confocal microscope (Zeiss LSM 510) with a colour digital camera. Images used for quantification were all single confocal sections. Photomicrographs were obtained with the following band-pass and long-pass filter settings: A488 (band pass filter: 505-530). The objective and the pinhole setting (1 airy unit, au) remained unchanged during the acquisition of a series for all images.

### Statistics

All data are presented as mean ± SEM. Statistical tests were performed with Prism 10 (GraphPad Software, La Jolla, CA, USA). No a priori sample size calculation was performed. Statistical analysis was undertaken only for studies where each group size was at least n=5.

The declared group size is the number of independent values, and statistical analysis was done using these independent values. Sample sizes were chosen (*i*) based on previous studies using similar methodologies and in line with standard practice in the field, and (*ii*) to comply with ethical considerations aimed at minimizing animal use, in accordance with the 3Rs principle, without compromising data quality. Normal distribution of data was analysed using the Shapiro-Wilk test. Depending on the experimental design, data were analysed using either Student’s t-test with equal variances, One-way ANOVA or Two-way ANOVA. The significance threshold was automatically set at p<0.05. ANOVA analyses were followed by Bonferroni *post hoc* test for specific comparisons only when overall ANOVA revealed a significant difference (at least p<0.05). Statistical details are reported in the **Suppl. Table 1**.

## Results

### NAc D2R-SPNs acquire responses to appetitive Pavlovian cues

D2R-SPNs respond to both food-related and food-predictive cues, as well as to food-related hormonal and nutrient signals, including triglycerides (TG) ^30,34,38–41^. Given their critical role in reward-related behaviours ^42^, we investigated whether this neuronal population (*i*) undergoes dynamic changes in activity during an appetitive Pavlovian conditioning and (*ii*) is modulated by circulating TG levels. To this end, we selectively expressed GCaMP6f in D2R-SPNs of *Drd2*-Cre mice (**Fig. 1A**) and monitored their Ca^2+^ activity throughout an appetitive Pavlovian conditioning paradigm (**Fig. 1B**).

**Figure 1:**
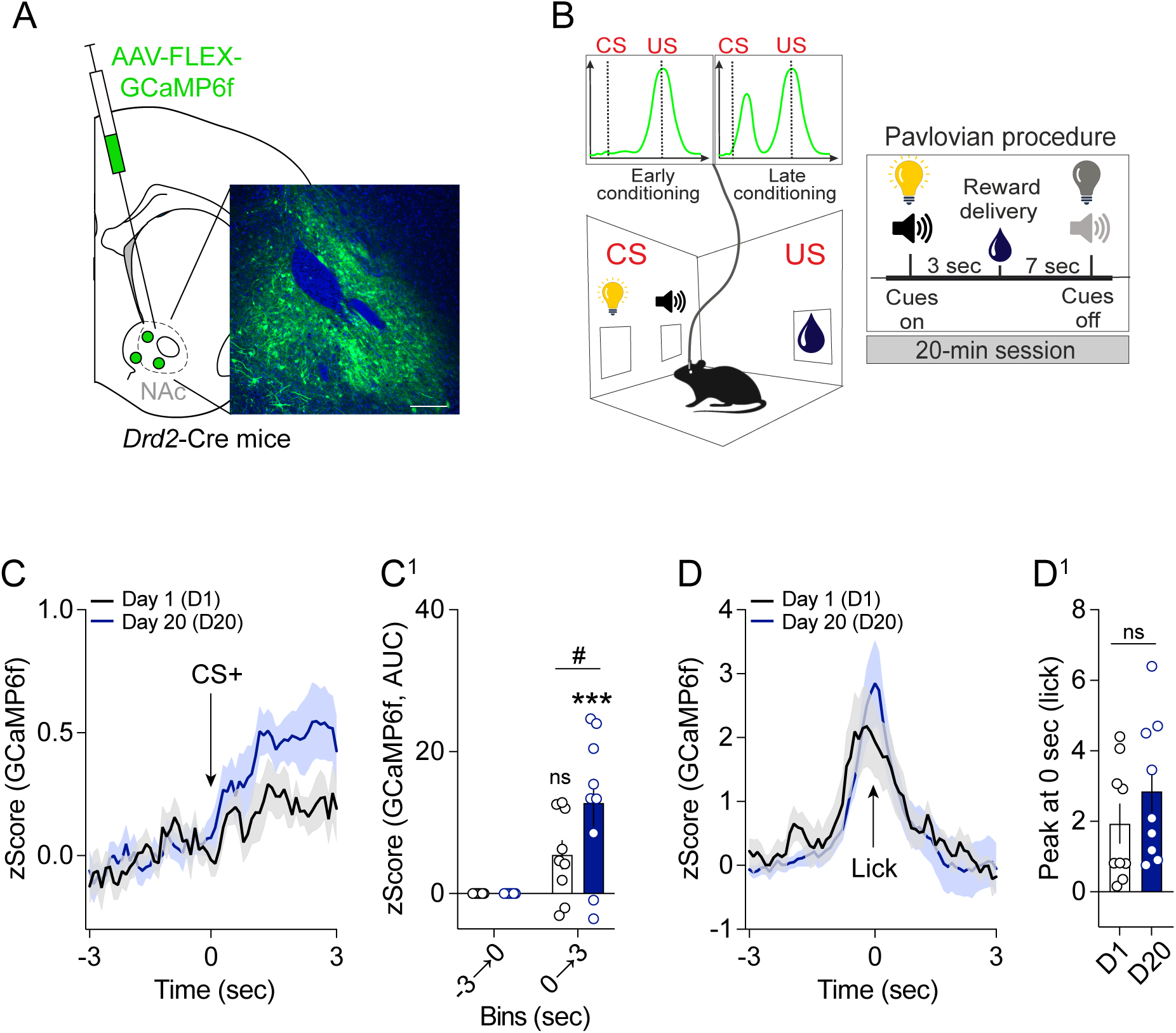
NAc D2R-SPNs respond to appetitive Pavlovian cues. (**A**) Detection of GCaMP6f in the NAc of *Drd2*-Cre mice. Scale bar: 200 μm. (**B**) Scheme representing the appetitive Pavlovian conditioning paradigm used in the study. (**C**) *In vivo* Ca^2+^ longitudinal dynamics of NAc D2R-SPNs and (**C^2^**) respective quantifications (AUC, area under the curb) during the presentation of CS^+^ in mice exposed to the operant chamber at Day 1 (D1, n=9) and at Day 20 (D20, n=9). (**D**) *In vivo* Ca^2+^ longitudinal dynamics of NAc D2R-SPNs and (**D^2^**) respective Ca^2+^ peak during the licking phase (food-reward consumption) in mice exposed to the operant chamber at Day 1 (D1, n=9) and at Day 20 (D20, n=9). Statistics: paired Student’s t-test (**C^2^, D^2^**), ***p<0.001 and ^#^p<0.05 for specific comparisons.

To validate the specificity of our recording approach, we first assessed D2R-SPN activity following systemic administration of amphetamine, a psychostimulant known to inhibit this neuronal population ^43^. As expected, amphetamine induced a significant reduction in D2R-SPN Ca^2+^ activity (**Suppl. Fig. 1A, A^2^**), confirming that our *in vivo* recordings reliably capture accumbal D2R-SPN activity.

During the first conditioning session (Day 1), D2R-SPNs did not exhibit a significant response to the conditioned stimulus (CS^+^; **Fig. 1C, C^1^**). In contrast, presentation of the food-reward elicited a robust increase in Ca^2+^ activity during consumption (lick; **Fig. 1D, D^1^**). Following repeated conditioning, D2R-SPNs progressively developed a strong response to CS^+^ presentation during the final conditioning session (Day 20; **Fig. 1C, C^1^**), indicating the acquisition of cue-evoked activity. Conversely, their response during reward consumption remained stable throughout training (**Fig. 1D, D^1^**).

Together, these findings demonstrate that accumbal D2R-SPNs are progressively recruited by reward-predictive cues during appetitive Pavlovian learning, while maintaining a stable response to reward consumption.

### Central TG availability modulates NAc D2R-SPN responsiveness in a metabolic state-dependent manner

We next investigated whether elevated central TG levels could modulate D2R-SPN responses to reward-predictive cues and food-reward consumption, and whether these effects depended on the nutritional state of the animals (food-restricted *versus* fed conditions). To selectively increase brain TG availability while avoiding peripheral alterations, TG or saline was delivered through an implanted carotid catheter ^2,25^.

In control mice maintained on standard chow diet (CD), central TG administration did not modify D2R-SPN responses to either CS^+^ presentation or reward consumption compared with saline-perfused controls (**Fig. 2A, A^1^, B, B^1^**). Furthermore, TG delivery in food-restricted mice did not alter reward-related behavioural parameters, including the number of licks or licking bursts (**Fig. 2C, D**).

**Figure 2:**
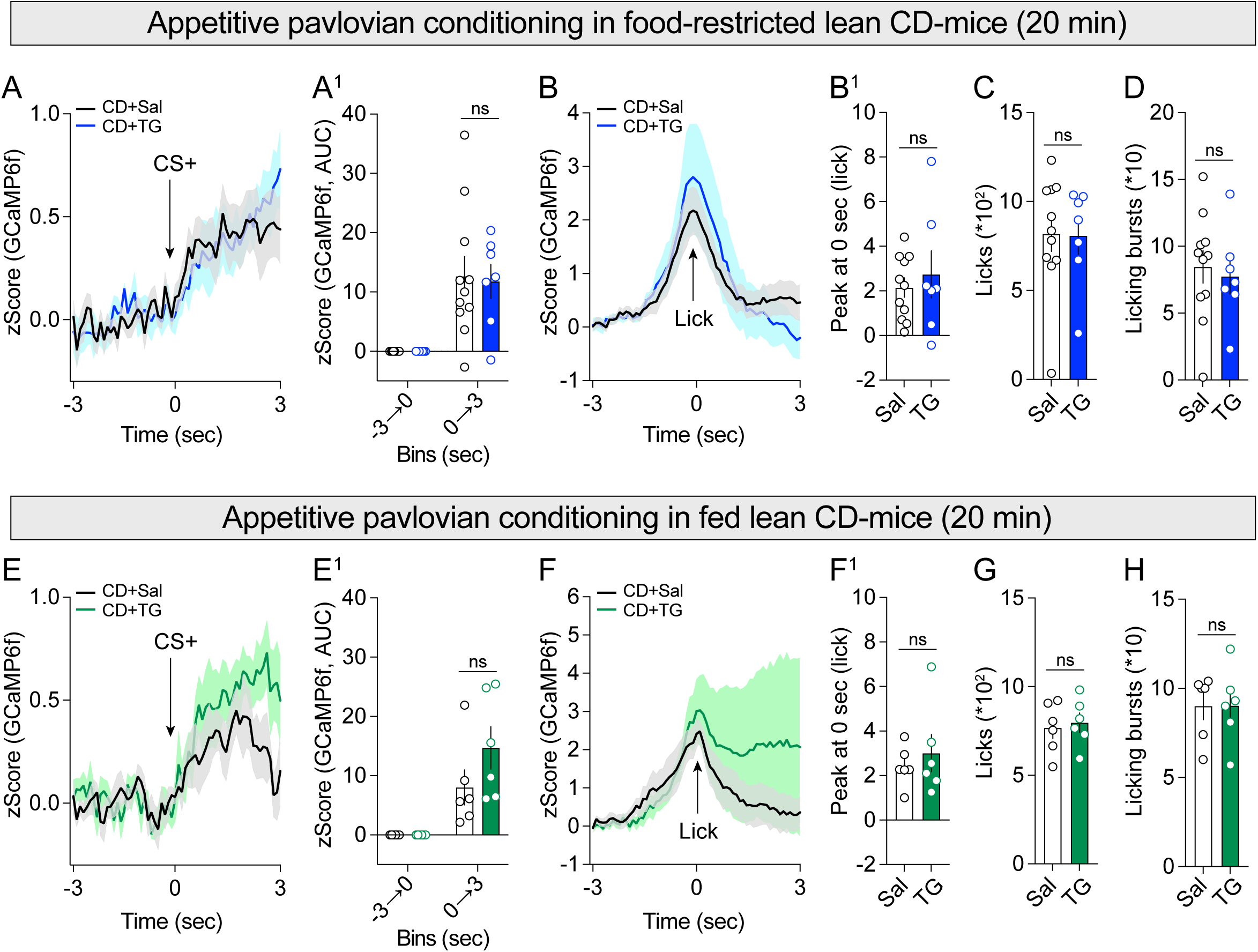
Central TG delivery does not alter D2R-SPN responses to appetitive Pavlovian cues in lean CD-mice. (**A**) *In vivo* Ca^2+^ longitudinal dynamics of NAc D2R-SPNs and (**A^1^**) respective quantifications (AUC, area under the curb) during the presentation of CS^+^ in food-restricted lean CD-mice receiving either saline (CD+Sal, n=11) or TG (CD+TG, n=7) and exposed to the Pavlovian operant chamber. (**B**) *In vivo* Ca^2+^ longitudinal dynamics of NAc D2R-SPNs and (**B^1^**) respective Ca^2+^ peak during the licking phase (food-reward consumption) in food-restricted lean CD-mice receiving either saline (CD+Sal, n=11) or TG (CD+TG, n=7) and exposed to the Pavlovian operant chamber. (**C**) Number of licks and (**D**) licking bursts in food-restricted lean CD-mice receiving either saline (CD+Sal, n=11) or TG (CD+TG, n=7) during the appetitive Pavlovian conditioning. (**E**) *In vivo* Ca^2+^ longitudinal dynamics of NAc D2R-SPNs and (**E^1^**) respective quantifications (AUC, area under the curb) during the presentation of CS^+^ in fed lean CD-mice receiving either saline (CD+Sal, n=6) or TG (CD+TG, n=6) and exposed to the Pavlovian operant chamber. (**F**) *In vivo* Ca^2+^ longitudinal dynamics of NAc D2R-SPNs and (**F^1^**) respective Ca^2+^ peak during the licking phase (food-reward consumption) in fed lean CD-mice receiving either saline (CD+Sal, n=6) or TG (CD+TG, n=6) and exposed to the Pavlovian operant chamber. (**G**) Number of licks and (**H**) licking bursts in food-restricted lean CD-mice receiving either saline (CD+Sal, n=6) or TG (CD+TG, n=6) during the appetitive Pavlovian conditioning. Statistics: Student’s t-test (**A^1^, B^1^, C, D, E^1^, F^1^, G, H**), ***p<0.001 and ^#^p<0.05 for specific comparisons.

We next assessed whether increased central TG availability could influence D2R-SPN responsiveness in fed mice undergoing appetitive Pavlovian conditioning. Similarly, central TG administration did not affect D2R-SPN Ca^2+^ activity during either CS^+^ presentation or food-reward consumption (**Fig. 2E, E^1^, F, F^1^**), nor did it alter licking behaviour or reward consumption parameters (**Fig. 2G, H**). Together, these results indicate that acute elevation of central TG levels does not influence D2R-SPN responses to reward-predictive cues or reward consumption under physiological metabolic conditions.

We next examined whether chronic exposure to an obesogenic diet (*e.g.*, high-fat diet, HFD) alters TG-mediated regulation of D2R-SPNs. Mice were exposed to a 1-month HFD before undergoing the pavlovian paradigm, and D2R-SPN responsiveness was assessed following central TG delivery in both food-restricted and fed states in HFD-mice.

Following food-restriction, central TG administration in HFD-fed mice resulted in a significant reduction of D2R-SPN Ca^2+^ activity in response to CS^+^ presentation compared with saline-perfused HFD controls (**Fig. 3A, A^1^**). In contrast, TG delivery did not modify D2R-SPN responses during reward consumption (**Fig. 3B, B^1^**). Importantly, these neuronal changes were not associated with alterations in behavioural performance, as licking responses and licking burst events remained unchanged (**Fig. 3C, D**).

**Figure 3:**
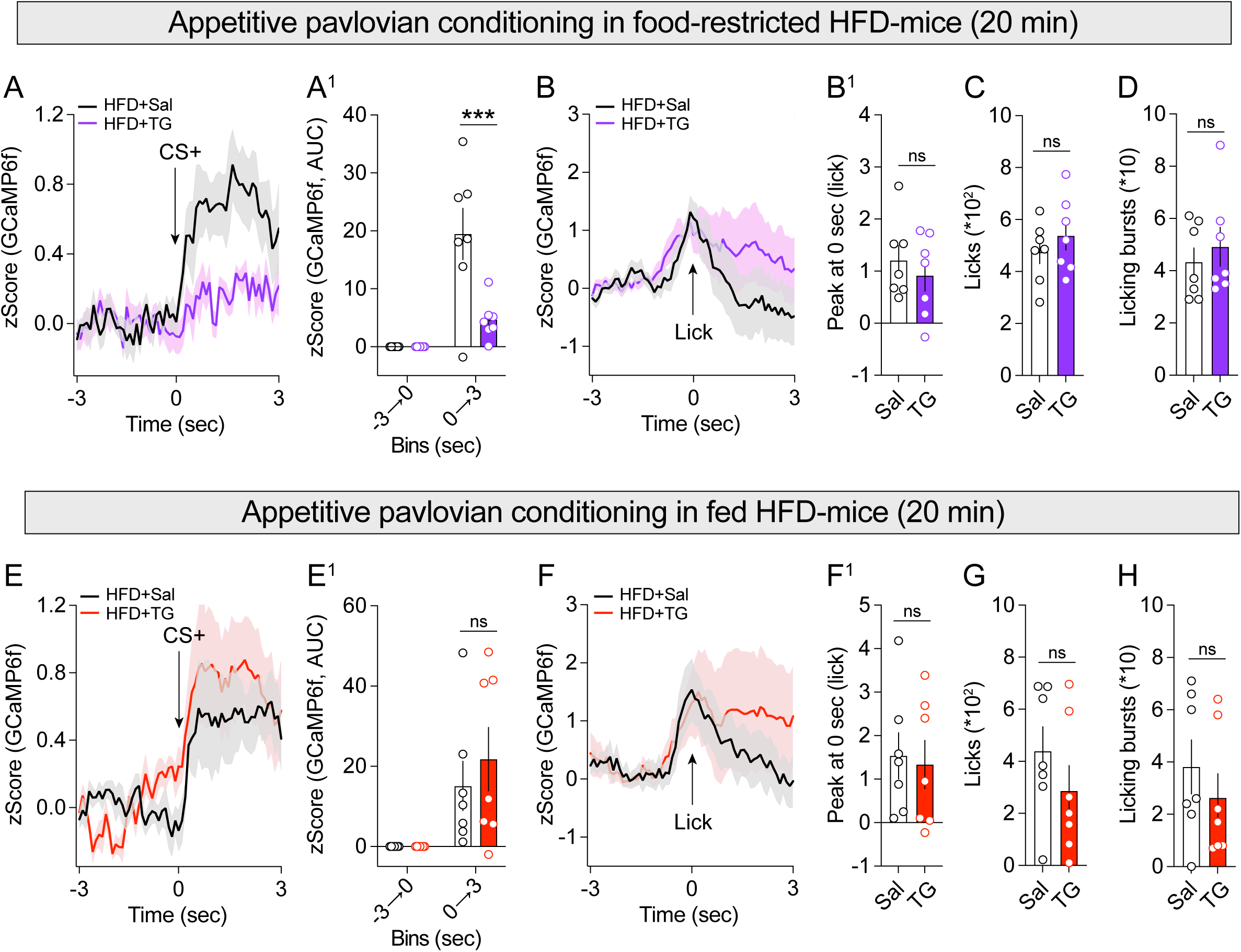
Central TG delivery alters D2R-SPN responses to appetitive Pavlovian cues in HFD-mice depending on the metabolic state. (**A**) *In vivo* Ca^2+^ longitudinal dynamics of NAc D2R-SPNs and (**A^1^**) respective quantifications (AUC, area under the curb) during the presentation of CS^+^ in food-restricted HFD-mice receiving either saline (HFD+Sal, n=7) or TG (HFD+TG, n=7) and exposed to the Pavlovian operant chamber. (**B**) *In vivo* Ca^2+^ longitudinal dynamics of NAc D2R-SPNs and (**B^1^**) respective Ca^2+^ peak during the licking phase (food-reward consumption) in food-restricted HFD-mice receiving either saline (HFD+Sal, n=7) or TG (HFD+TG, n=7) and exposed to the Pavlovian operant chamber. (**C**) Number of licks and (**D**) licking bursts in food-restricted HFD-mice receiving either saline (HFD+Sal, n=7) or TG (HFD+TG, n=7) during the appetitive Pavlovian conditioning. (**E**) *In vivo* Ca^2+^ longitudinal dynamics of NAc D2R-SPNs and (**E^1^**) respective quantifications (AUC, area under the curb) during the presentation of CS^+^ in fed HFD-mice receiving either saline (HFD+Sal, n=7) or TG (HFD+TG, n=7) and exposed to the Pavlovian operant chamber. (**F**) *In vivo* Ca^2+^ longitudinal dynamics of NAc D2R-SPNs and (**F^1^**) respective Ca^2+^ peak during the licking phase (food-reward consumption) in fed HFD-mice receiving either saline (HFD+Sal, n=7) or TG (HFD+TG, n=7) and exposed to the Pavlovian operant chamber. (**G**) Number of licks and (**H**) licking bursts in fed HFD-mice receiving either saline (HFD+Sal, n=7) or TG (HFD+TG, n=7) during the appetitive Pavlovian conditioning. Statistics: Student’s t-test (**A^1^, B^1^, C, D, E^1^, F^1^, G, H**), ***p<0.001 for specific comparisons.

Interestingly, central TG delivery in fed HFD-mice did not affect D2R-SPN responses to either CS^+^ presentation or reward consumption, nor did it alter reward-related behavioural parameters (**Fig. 3E-H**).

Together, these findings demonstrate that obesogenic conditions unmask a TG-sensitive state in D2R-SPNs, characterized by a selective reduction of cue-evoked activity in food-restricted animals. This effect depends on metabolic state, suggesting that HFD exposure alters the sensitivity of D2R-SPNs to metabolic modulation by TG.

### Central TG delivery in HFD-mice reduces NAc D2R-SPN activity in response to refeeding

Given the altered D2R-SPN responsiveness observed following TG administration in HFD-mice during Pavlovian conditioning, we next investigated whether TG could also modulate D2R-SPN activity during natural food consumption.

In overnight-fasted CD-mice, central TG delivery did not affect D2R-SPN Ca^2+^ activity in response to refeeding (**Fig. 4A, A^1^**), nor did it alter fasting-induced food intake (**Fig. 4B**). In contrast, in overnight-fasted HFD-mice, central TG administration significantly reduced refeeding-evoked D2R-SPN Ca^2+^ activity (**Fig. 4C, C^1^**), while food intake remained unchanged (**Fig. 4D**).

**Figure 4:**
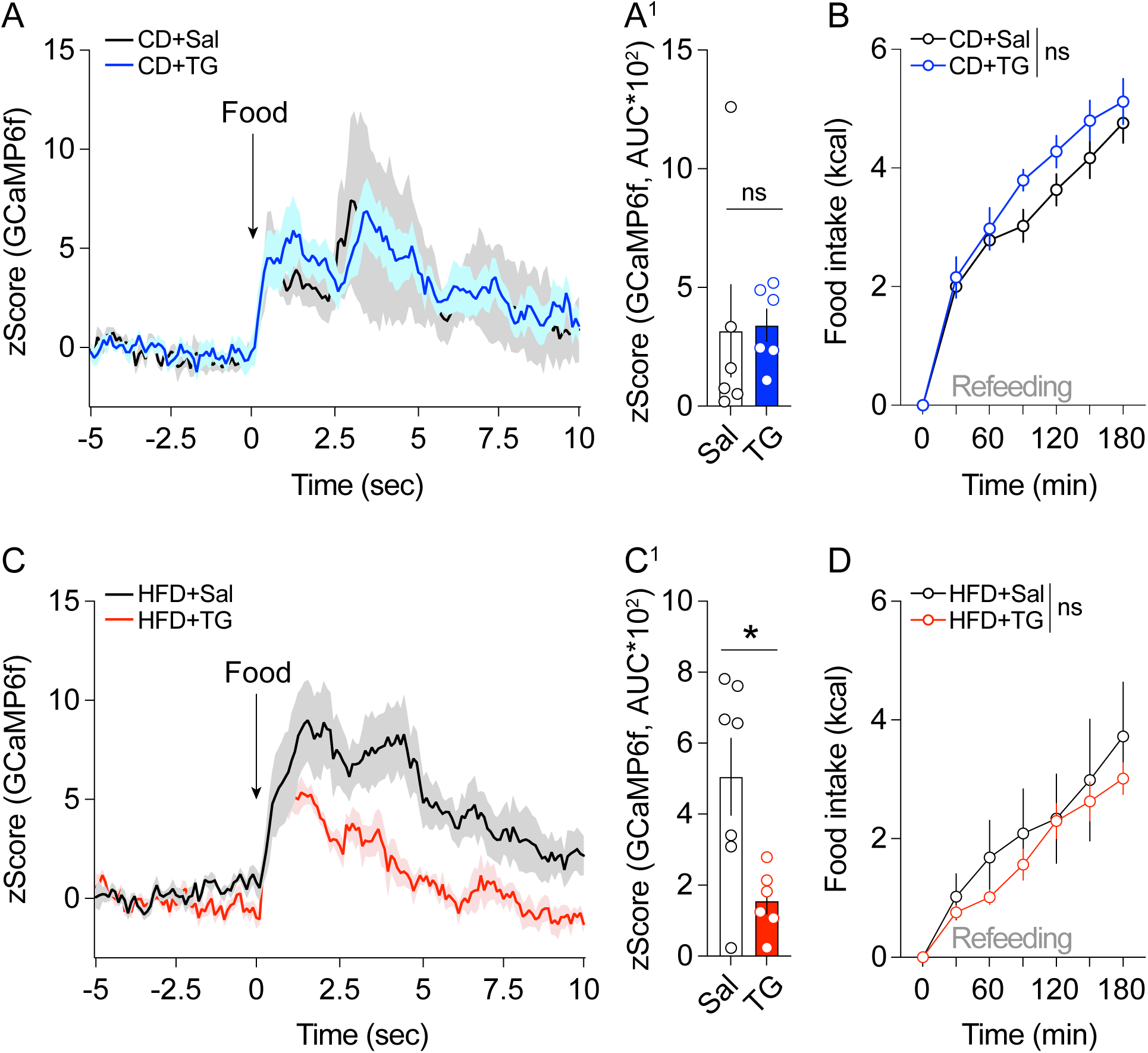
Central TG delivery alters D2R-SPN responses to refeeding in HFD-mice but not CD-mice. (**A**) *In vivo* Ca^2+^ longitudinal dynamics of NAc D2R-SPNs and (**A^1^**) respective quantifications (AUC, area under the curb) during refeeding on CD in food-deprived lean CD-mice receiving either saline (CD+Sal, n=6) or TG (CD+TG, n=6). (**B**) Food intake (kcal) during the refeeding paradigm. (**C**) *In vivo* Ca^2+^ longitudinal dynamics of NAc D2R-SPNs and (**C^1^**) respective quantifications (AUC, area under the curb) during refeeding on HFD in food-deprived HFD-mice receiving either saline (HFD+Sal, n=7) or TG (HFD+TG, n=6). (**D**) Food intake (kcal) during the refeeding paradigm. Statistics: Student’s t-test (**A^1^, C^1^**), *p<0.05 for specific comparisons, and Two-way ANOVA (**B, D**).

These findings indicate that obesogenic conditions increase the sensitivity of D2R-SPNs to TG-mediated regulation during energy restoration, leading to reduced neuronal activity during food consumption without affecting the amount of food ingested.

### Central TG delivery does not affect NAc D2R-SPN responses during novelty exploration

Finally, we investigated whether TG-mediated suppression of D2R-SPN activity in HFD-mice represented a generalized alteration in neuronal responsiveness or was restricted to food- and reward-related contexts.

To this end, mice were exposed to a novel environment, a behavioural paradigm known to transiently recruit D2R-SPNs ^44^. Central TG delivery did not modify novelty-induced D2R-SPN Ca^2+^ dynamics compared with saline controls, irrespective of metabolic status (CD *versus* HFD conditions; **Fig. 5A-B^1^**).

**Figure 5:**
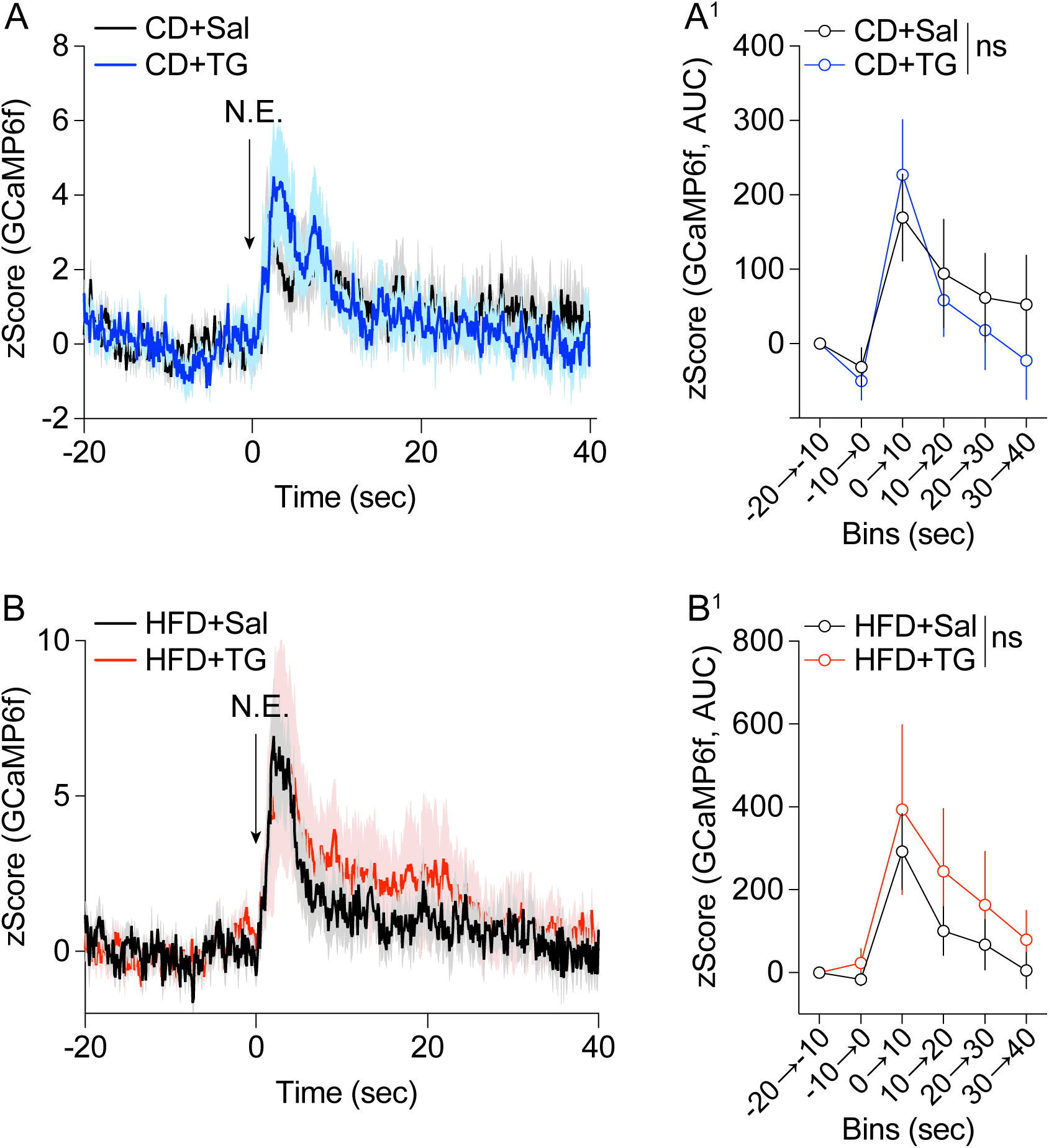
Central TG delivery does not alter D2R-SPN responses to a new environment. (**A**) *In vivo* Ca^2+^ longitudinal dynamics of NAc D2R-SPNs and (**A^1^**) respective quantifications (AUC, area under the curb) during the exploration of a new environment in lean CD-mice receiving either saline (CD+Sal, n=6) or TG (CD+TG, n=6). (**B**) *In vivo* Ca^2+^ longitudinal dynamics of NAc D2R-SPNs and (**A^1^**) respective quantifications (AUC, area under the curb) during the exploration of a new environment in HFD-mice receiving either saline (HFD+Sal, n=8) or TG (HFD+TG, n=8). Statistics: Two-way ANOVA (**A^1^, B^1^**).

Together, these results indicate that TG-mediated modulation of D2R-SPN activity is not due to a generalized reduction in neuronal excitability but instead reflects a context- and metabolic state-dependent regulation of D2R-SPN function, selectively emerging during food- and reward-related behaviours under obesogenic conditions.

## Discussion

The present study provides the first direct evidence that circulating triglycerides (TG) modulate accumbal D2R-SPN activity during appetitive behaviours in a metabolic state-dependent manner. Using *in vivo* fiber photometry in *Drd2*-Cre mice, we demonstrate that central TG delivery selectively suppresses cue-evoked D2R-SPN responses in food-restricted mice maintained on a high-fat diet (HFD), but not in chow-diet (CD) mice under any metabolic state. These findings reveal a novel mechanism by which dietary lipids interface with the accumbal reward circuit to regulate motivated behaviours, with important implications for understanding how obesogenic diets alter neural processing of food-related cues.

Our observation that D2R-SPNs progressively acquire CS^+^ responses during Pavlovian conditioning extends previous work on accumbal DA signaling during associative learning ^45–47^. While DA release in the NAc is well-established to encode reward prediction errors and cue-reward associations ^48,49^, the specific contributions of D2R-SPN populations to this process have remained less clear. Recent studies have revealed dynamic shifts in D2R-SPN activity during learning and consummatory behaviours ^34^, with D2R-SPNs showing sustained activity during food approach and decreased activity upon consumption onset ^34,50,51^. Moreover, while chemogenetic manipulation of NAc D1R-SPNs enhanced motivation for food reward and voluntary exercise and reduced food intake, chemogenetic activation of D2R-SPNs produced opposite effects ^34^.

Our findings align with this emerging view that D2R-SPNs are not simply inhibitory to reward processes, but rather encode specific aspects of motivated behaviours in a context-dependent manner.

The progressive acquisition of CS^+^ responses we observed in D2R-SPNs contrasts with the traditional view that striatopallidal neurons primarily inhibit reward-related behaviours ^52,53^. Instead, our data support a more nuanced model in which D2R-SPNs encode learned associations between predictive cues and rewarding outcomes. This is consistent with recent optogenetic and chemogenetic studies showing that D2R-SPN activation can enhance high-fat food intake in specific contexts, and that NAc D1R- and D2R-SPNs have distinct, valence-independent roles in learning ^51,54^, as well as experience-dependent encoding of “liking” and “wanting” components of hedonic feeding ^50^. The fact that D2R-SPN responses to the CS^+^ increased across training sessions suggests these neurons participate in the encoding of incentive salience, potentially by modulating the balance between approach and avoidance behaviours during goal-directed actions ^39^.

The most striking finding of our study is that TG delivery selectively reduces cue-evoked D2R-SPN activity in HFD-mice under food restriction, but has no effect in CD-mice regardless of metabolic states. This observation provides evidence supporting the hypothesis that circulating lipids gate DA-associated behaviours through D2R-expressing neurons ^2^. In fact, we have previously demonstrated that nutritional TG can be metabolized within the mesocorticolimbic system via the lipoprotein lipase (LPL) and modulate D2R-SPN activity ^2^. The present study extends this work by showing that the sensitivity of D2R-SPNs to TG signaling is not constitutive, but rather depends on both prior dietary history (HFD vs. CD) and current metabolic states (food-restricted vs. fed).

The fact that intracarotid TG infusion decreased NAc D2R-SPN activity only in food-restricted mice in obesogenic condition suggests a form of metabolic priming. Chronic HFD consumption is known to induce numerous adaptations within the mesolimbic DA system, including downregulation of striatal D2Rs ^14,15,30,55–58^, altered DA release and reuptake dynamics ^12,30^, and changes in synaptic transmission regulating DA release in the NAc ^11,13,15,30,56,59^. These HFD-induced neuroadaptations may render D2R-SPNs more sensitive to circulating metabolic signals, including TG. Indeed, obesity and HFD exposure have been shown to disrupt DA neurotransmission through convergent effects of metabolic state, physiological stress, and inflammation on the dopaminergic control of food intake ^60^.

The additional requirement for food restriction to unmask TG effects on D2R-SPNs is particularly intriguing. Food restriction and fasting are known to profoundly alter accumbal DA signaling, with metabolic hormones such as ghrelin acting as an interface between physiological state and phasic DA responses ^61^. Ghrelin increases the magnitude of food-evoked DA spikes through a mechanism involving lateral hypothalamic orexin-neurons that project to the VTA ^61^ and hypothalamic-accumbal circuits may relay information about circulating TG levels to modulate striatal activity in a state-dependent manner.

Our findings suggest that the fasted state may similarly sensitize D2R-SPNs to metabolic signals, potentially through altered expression or function of lipid-sensing machinery. While primarily described in the hypothalamus, a well-studied site of lipid sensing (for review ^4,7,62^), metabolic regulation of TG-processing enzymes, including but not limited to LPL, have been documented ^4,63–68^. In the NAc, such modulation of lipase expression, function and co-activators availability could also be at play to modulate local TG availability. In addition, brain trafficking of TG-containing lipoprotein could also modulate local TG availability in the brain in a metabolic-dependent manner ^63,64^. Finally, lipid droplets and TG deposit in neurons have only recently been documented as integral components of synaptic transmission ^3,5^ and could also serve as signaling mechanisms mediating metabolic priming of brain circuits following high-fat feeding, ultimately shaping the acute response of the brain to postprandial increases in TG.

A key observation in our study is that TG delivery specifically reduced CS^+^-evoked D2R-SPN activity in food-restricted HFD-mice, while having no effect on novelty-induced D2R-SPN activity in any condition. This specificity suggests that TG signaling does not globally suppress D2R-SPN excitability, but rather selectively modulates responses to learned food-predictive cues.

This cue-specific modulation may reflect the integration of metabolic information with learned associations in the NAc. The NAc is positioned at the interface between limbic and motor systems, integrating information about motivational state, learned associations, and action selection ^69^. D2R-SPNs, which project primarily to the ventral pallidum via the indirect pathway, are thought to regulate motivation by modulating inhibitory transmission ^70^. Upregulation of D2R in NAc D2R-SPNs has been shown to increase willingness to work for food by decreasing inhibitory transmission to the ventral pallidum ^70^. Our findings suggest that TG signaling may act as a satiety-like signal that reduces the motivational impact of food-cues by dampening D2R-SPN responses, but only when the system has been primed by chronic HFD exposure and is in a metabolically vulnerable states (*e.g.*, food restriction).

### Brain TG sensing and D2R-SPN responses to refeeding

We also observed that TG delivery oppose D2R-SPN responses to food cues in fasted HFD-mice. This finding extends the cue-specific effects to consummatory behaviours and suggests that TG signaling may serve as a negative feedback signal. The NAc has been implicated in the control of feeding and energy homeostasis, with specific D1R- and D2R-expressing neuronal subtypes playing distinct roles ^71,72^. Recent studies identified a subset of DA receptor-expressing neurons in the NAc shell that controls feeding and energy homeostasis, and demonstrated that a putative loop connection between VTA DA-neurons and NAc encodes positive valence to compensate for hunger ^71,72^

The reduction in refeeding-induced D2R-SPN activity by TG delivery in HFD fasted mice may reflect a mechanism by which post-prandial increased in plasma lipids signal nutrient availability to reduce the drive to consume food. This is consistent with the broader literature on lipid sensing in the hypothalamus, where fatty acids and their metabolites regulate feeding behaviours and energy homeostasis (for review ^4,62^). The fact that this effect was only observed in HFD-mice again highlights the importance of dietary history in determining sensitivity to metabolic signals. Chronic HFD exposure may alter the expression and/or function of lipid-sensing receptors or downstream signaling pathways in D2R-SPNs or their afferent inputs, rendering these neurons more responsive to TG fluctuations.

Our findings have important implications for understanding the neural mechanisms underlying obesity and compulsive eating. Downregulation of striatal D2 receptors is a hallmark of obesity in both humans and rodents ^57,73,74^, and knockdown of striatal D2 receptors accelerates the development of addiction-like reward deficits and compulsive-like food seeking in rats with extended access to palatable high-fat food ^58^. The observation that TG signaling selectively modulates D2R-SPN activity in HFD-mice suggests that altered lipid metabolism may contribute to reward dysfunction in obesity.

Interestingly, the inhibitory action of TG onto D2R-SPN activity, might initially seem counterintuitive given that D2R-SPNs are thought to inhibit reward processes. However, recent work has challenged this simple dichotomy, showing that D2R-SPNs can promote consumption under certain conditions. In the context of hedonic feeding, activation of D2R-SPNs has been shown to enhance high-fat food intake, while their inhibition reduces intake ^34,54^. Our finding that TG delivery reduces D2R-SPN activity in HFD food-restricted mice may therefore represent a compensatory mechanism to limit excessive food intake in response to elevated circulating lipids. The failure of this mechanism in the fed state, where TG had no effect, may contribute to overconsumption and weight gain. In condition of chronic hypertriglyceridemia associated with obesity, one may hypothesis that a maladaptive desensitization occurs, resulting in altered D2R-SPN response to acute post-prandial TG.

The metabolic regulation of TG signalling also has implications for understanding individual differences in obesity susceptibility. Obesity-prone and obesity-resistant rats show differential responses to palatable food in terms of NAc dopamine receptor expression, with D1R expression decreasing in obesity-prone animals after long-term exposure to palatable food ^75,76^. Our findings suggest that sensitivity to circulating TG signals may similarly differ between individuals, potentially contributing to differences in the ability to regulate food intake in response to metabolic cues. Future studies examining TG effects on D2R-SPN activity in obesity-prone versus obesity-resistant animals could provide important insights into the neural basis of obesity susceptibility.

Some limitations of the current study should be acknowledged. First, we used central (carotid catheter) TG delivery to manipulate circulating TG levels, which may not fully recapitulate the physiological dynamics of postprandial TG excursions. Future studies using dietary manipulations to induce more naturalistic TG fluctuations would be valuable. Second, we focused on D2R-SPNs in the NAc, but did not distinguish between core and shell subregions, which are known to have distinct functional roles. Third, while we observed clear effects of TG on D2R-SPN activity, we did not measure the activity of D1R-SPNs.

## Conclusions

In summary, our study demonstrates that circulating TG modulate accumbal D2R-SPN responses to learned food cues in a metabolic state-dependent manner. This modulation requires both chronic HFD exposure and food restriction, suggesting a form of metabolic priming that renders D2R-SPNs sensitive to TG signaling. The specificity of TG effects for learned cue responses, but not novelty-induced activity, indicates that TG signaling selectively modulates the incentive salience of food-predictive cues rather than affecting general arousal or novelty processing. These findings provide novel insights into how dietary lipids interface with striatal reward circuits to regulate motivated behaviours, and have important implications for understanding the neural mechanisms underlying obesity and compulsive eating. Future studies elucidating the molecular and circuit mechanisms of TG signaling in the NAc may identify new therapeutic targets for obesity and related metabolic disorders.

## Acknowledgments

We thank Olja Kacanski for administrative support; Isabelle Le Parco, Daniel Quintas, Magguy Boa, Ludovic Maingault, Angélique Dauvin, Siscaro Léa and Florianne Michel for animals’ care. We also thank Dr. Claire Martin, Dr. Federica Genovese, Dr. Maya Faour and Dr. Oriane Onimus for discussions, help and assistance. We acknowledge the Functional and Physiological Exploration platform (FPE) of Université Paris Cité, CNRS, *Unité de Biologie Fonctionnelle et Adaptative*, and the animal core facility “Buffon” of the Université Paris Cité/Institut Jacques Monod.

## Author contributions

Guangping Li: Investigation; Methodology; Visualization; Formal analysis; Data curation; Software.

Julien Castel: Investigation; Methodology.

Anthony Ansoult: Methodology.

Benoit Bertrand: Methodology.

Serge Luquet: Conceptualization; Investigation; Funding acquisition; Writing - original draft; Methodology; Validation; Visualization; Writing - review & editing; Formal analysis; Project administration; Software; Data curation; Supervision; Resources.

Giuseppe Gangarossa: Conceptualization; Investigation; Funding acquisition; Writing - original draft; Methodology; Validation; Visualization; Writing - review & editing; Formal analysis; Project administration; Software; Data curation; Supervision; Resources.

## Data availability

The data that support the findings of this study are available from the corresponding author upon reasonable request.

---

**Suppl. Figure 1: Amphetamine administration inhibits D2R-SPN activity.** (**A**) *In vivo* Ca^2+^ longitudinal dynamics of NAc D2R-SPNs and (**A^1^**) respective quantifications (AUC, area under the curb) following the administration of amphetamine (2 mg/kg, i.p., n=7). Statistics: paired Student’s t-test (**A^1^**), *p<0.05 for specific comparisons.

