## Supplementary figures and images for "Central lipid sensing tunes accumbal D2R-neuron activity in appetitive behaviours"

### Suppl. Figure 1

#### Suppl. Figure 1

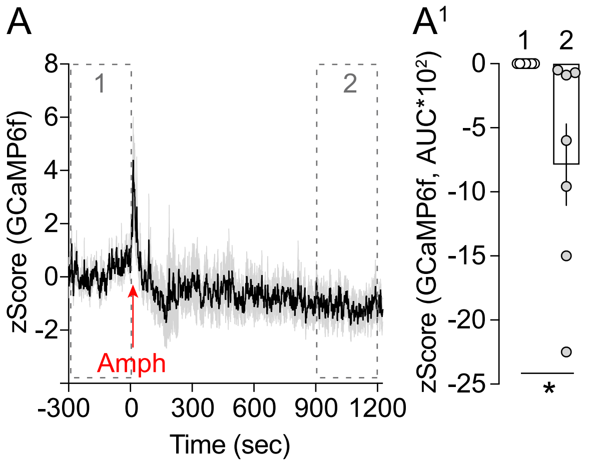
